# Rapid high-resolution measurement of DNA replication timing by droplet digital PCR

**DOI:** 10.1101/208546

**Authors:** Dzmitry G. Batrakou, Emma D. Heron, Conrad A. Nieduszynski

## Abstract

Genomes are replicated in a reproducible temporal pattern. Current methods for assaying allele replication timing are time consuming and/or expensive. These include high-throughput sequencing which can be used to measure DNA copy number as a proxy for allele replication timing. Here, we use droplet digital PCR to study DNA replication timing at multiple loci in budding yeast and human cells. We establish that the method has temporal and spatial resolutions comparable to the high-throughput sequencing approaches, while being faster than alternative locus-specific methods. Furthermore, the approach is capable of allele discrimination. We apply this method to determine relative replication timing across timing transition zones in cultured human cells. Finally, multiple samples can be analysed in parallel, allowing us to rapidly screen kinetochore mutants for perturbation to centromere replication timing. Therefore, this approach is well suited to the study of locus-specific replication and the screening of *cis-* and *trans-acting* mutants to identify mechanisms that regulate local genome replication timing.

## INTRODUCTION

DNA replication is the process by which genetic information is duplicated before transmission from parental to daughter cells. Quantification of DNA replication is, therefore, a measurement of DNA within the biologically restricted scale of between one and two. At a single molecule level, a region of DNA is either unreplicated and has a copy number of one, or has been replicated and has a copy number of two. However, in ensemble samples, differences in cell cycle synchrony and the stochastic nature of DNA replication lead to variations in the proportions of cells that have a particular locus replicated. Thus, instead of the discrete binary scale that can be applied to a single molecule, average DNA copy number per cell in a synchronised population lies on a continuous scale from one to two. This relative copy number is inversely proportional to the locus’ replication timing: it is close to two in early replicated regions of DNA and approaching a relative copy number of one in late, passively replicated regions. Therefore relative copy number of DNA from a replicating cell population can serve as a proxy for DNA replication timing (1).

Bacteria, such as *Escherichia coli,* have small genomes with a single origin of replication per replication unit (chromosome or plasmid) from which DNA replication is initiated. Eukaryotes have much larger genomes and contain multiple origins per replication unit, presumably to facilitate the timely completion of S phase and protect the stability of large replication units (2–5). In this case, origins fire with different efficiencies (proportion of cells in a population in which the origin is active) and timing (relative to the onset of S phase). Notably, the timing of origin firing follows a defined and reproducible replication programme in many organisms and tissues (6–8). This phenomenon is poorly understood. However, it is known that some human disorders involve changes in the replication timing programme, for example, in many types of cancer (9). As yet, it is unclear whether those changes are a cause or consequence of these disorders.

Methods used to study DNA replication can be divided by the scope of investigation: locus studies that use specific probes or genome-wide approaches that take advantage of microarrays or high-throughput sequencing (HTS). In addition, some methods use direct DNA copy number quantification, for example locus-specific microscopy-based approaches (10, 11), quantitative real-time PCR (qPCR) (12) or deep sequencing (1, 13). Other methods rely on the incorporation of labelled bases into the nascent strand during DNA replication (14–16).

The original nascent-strand based method for analysis of DNA replication timing involves the incorporation of bases with C and/or N substituted for dense isotopes in an adaption of Meselson and Stahl’s famous experiment in support of semiconservative DNA replication. The dense isotope transfer assay, as first established, was a single-locus, quantitative assay (17) that has more recently been applied genome-wide (18). However, it is time-consuming, requires defined media for cell culture and the variable density of genomic fragments can result in artefacts (19). An alternative to dense isotopes involves the incorporation of base analogues at actively replicating forks, which allows subsequent immunoprecipitation of nascent DNA for analysis by qPCR, microarrays or HTS. This is the basis for repli-seq, which involves sorting subfractions of S phase cells after a pulse of labelling with a thymidine analogue and HTS of immunoprecipitated nascent DNA (14, 16, 20). Sequences that are enriched in the early S phase population are defined as early replicating, while sequences enriched in the late S phase sample are those that replicate late. This method can detect transition zones between early and late replicating regions of genomic DNA.

The microscopy-based replication timing methods use a single-cell, locus-specific copy number approach. The locus of choice is labelled either by FISH or targeted by a fluorescent protein (facilitated by the tetO-TetR interaction, or similar) and its replication detected by the duplication of the labelled loci (10, 11). This method relies on cell synchronisation and requires involved sample processing (or genome editing, when labelling with fluorescent proteins), data acquisition and analysis. In the case of mammalian cells, the need for cell synchronisation may be mitigated by the simultaneous visualisation of replication sites and binning replication timing into 5 phases with distinct patterns (21). Nuclease-dead Cas9 has been utilised to avoid complicated sample processing or genome engineering (22) while allowing live imaging (23). However, the method is likely to remain of low throughput.

Other copy number methods that determine replication timing rely on measuring relative DNA copy number either in asynchronous cell cultures or S phase samples (1, 24). The latter can be obtained by sorting using fluorescence activated cell sorting (FACS) to isolate the whole S-phase cell population, or cells can be synchronised in S phase. Deep sequencing can then be used to precisely analyse relative DNA copy number from S phase cells. We have previously shown mathematically and experimentally that relative DNA copy number in S phase cells is linearly related to the relative replication time (1, 25). While this HTS-based approach is valuable for the measurement of genome-wide replication timing, the temporal precision and spatial resolution depend on sequencing depth, and is less practical for organisms with large genomes, such as humans, and in studies where high spatial resolution is required. For single allele studies, quantitative real-time PCR (qPCR) offers the benefit of a cheap, fast and high throughput approach that has been used to detect differences in DNA replication time (12). However, it requires a large number of technical replicates to detect small differences and the detection limit can be greatly affected by impurities in DNA samples (26, 27). Therefore, there is a need for a more precise, locus-specific, quantitative method to study DNA replication that is also less sensitive to contaminants.

Digital PCR is a recent refinement of quantitative PCR (28–31). It relies on partitioning of a qPCR mix into many reactions each of much smaller volume. In these partitioned reactions, all components are present in vast excess, except for the target template DNA molecules. For example, droplet digital PCR takes advantage of non-miscible liquid phases to generate compartments of around 1 nL in size (31). This allows easy handling of the digital PCR reactions in widely used 96-well format and bypasses the requirement for expensive microfluidics devices making it the cheapest, per sample, digital PCR platform currently available. For successful quantification in digital PCR, DNA samples must be diluted so that only a fraction of the partitions contain the target DNA. Following end-point PCR amplification and fluorescence detection, partitions are either positive or negative, thus resulting in a digital readout. The proportion of positive partitions is then used to calculate the number of target DNA molecules in the initial PCR mix. Therefore, the resulting method provides absolute quantification of target DNA in a single reaction without the need for the calibrating standard curves used in traditional qPCR. Compared to qPCR, digital PCR offers higher precision and is less sensitive to DNA sample contamination (32). The method has already been used successfully to compare replication timing between SNP-containing alleles in human lymphoblastoid cell lines (8).

Here, we confirm that droplet digital PCR (ddPCR) can be used to measure relative DNA copy number during DNA replication. Using *Saccharomyces cerevisiae* and human cell lines, we compare ddPCR to HTS-based replication timing analyses, testing the temporal and spatial resolution, and the ability to distinguish alleles. We demonstrate that this method can be applied to organisms with large genomes by quantifying the relative replication timing across timing transition zones in cultured human cells. Finally, we find that the throughput of this method allows for the rapid screening of multiple mutants to determine locus-specific perturbations to replication timing.

## MATERIAL AND METHODS

### Strains and cell lines

Yeast strains used in this study are listed in Supplementary Table 1. Cells were grown in standard rich YPAD medium (Formedium). HeLa and Jurkat cells were cultured in DMEM supplemented with 10% v/v FBS and 100 u/mL each Penecillin and Streptomycin (Gibco) at 37°C in a humid atmosphere with 5% CO^2^. MRC-5 cells were cultured as above but supplemented with 20% v/v FBS. To arrest MRC-5 cells in G1 phase, cells were cultured in medium lacking FBS for 7 days (with one medium change after 3 days).

### Time course experiments

For cell cycle synchronisation, yeast cells were grown, arrested and released at 23°C. Alpha factor was added at OD^600^ ~0.2 to a final concentration of 450 nM with subsequent additions to maintain the arrest for 1.5-2 generation times; release was initiated by addition of pronase to 0.2 mg/ml (zero time point). Culture samples were collected at the indicated times and immediately mixed with 10% volume of ice-cold AE buffer (1% sodium azide, 0.2 M EDTA pH 8.0) for flow cytometry analysis and DNA extraction. All cells were pelleted by centrifugation and washed once with water. For DNA extraction, cell pellets were stored at -20°C. For flow cytometry analysis, cells were fixed in 70% ethanol for a minimum of 10 hours at 4°C.

### Flow cytometry and cell sorting

#### Yeast

Cells were grown at 30°C to an OD^600^ of 0.5–0.7. Cells were pelleted, washed twice with water and fixed in 70% ethanol for a minimum of 10 hours at 4°C. Cells may be stored long term during this step. Fixed cells were pelleted, washed twice, resuspended in FC buffer (50 mM sodium citrate pH7.0, 0.1% sodium azide), and treated consecutively with 0.1 mg/ml RNase A and 0.2 mg/ml proteinase K, for 1 hour each at 55°C. To stain DNA, the cells were resuspended in FC buffer containing 2 μM (flow cytometry) or 10 μM (FACS) SYTOX™ Green Nucleic Acid stain (Invitrogen) and incubated overnight at 4°C. Prior to analysis, cells were pulse sonicated to break cell clumps and diluted two-fold with FC buffer. Flow cytometry samples were analysed on a Cytek DxP flow cytometer using the 488 nm laser and 530/30 filter. A MoFlo Sorter (Coulter Beckman) was used to sort 1-5 million cells from respective cell cycle stages. The DNA fluorescence histogram plot was used to set the gates for the sorting. The purity of the sorted cell fractions was confirmed by flow cytometry.

#### Human cells

Cells were washed once in PBS and trypsinised. Trypsin was neutralised with PBSF (PBS supplemented with 2% FBS) and the cells were pelleted, rinsed twice in PBSF, resuspended in PBS and fixed in 70% ethanol for at least 1 h at 4°C. Cells may be stored long term during this step. After fixing, cells were rinsed twice with PBSF and incubated in staining solution (PBSF supplemented with 3.8 mM sodium citrate, 5 μg/ml RNase A and 50 μg/ml of Propidium Iodide, PI) for 30 min at ambient temperature in the dark. Alternatively, 2 μM SYTOX™ Green Nucleic Acid stain may be used in place of Propidium Iodide followed by overnight incubation at 4°C (as was done in the experiment in Supplemental Figure 7). An Astrios (Beckman Coulter) sorter was used to sort partial G1 and whole S phase fractions (at least 3 million each) from HeLa cells based on PI signal.

### DNA extraction

#### Yeast genomic DNA

Sorted cells were flocculated by addition of ethanol to 30% v/v final and pelleted. Cell pellets (either sorted or from time course) were resuspended in 50 mM Tris-HCl pH 8.0, 0.1 M EDTA, 0.1% v/v β-mercapthoethanol and were spheroplasted with 1 mg/ml Zymolyase for 30 min at 37°C. The reactions were supplemented with 1% w/v SDS, 0.2 M NaCl, 0.1 mg/ml RNase A and 0.2 mg/ml proteinase K and incubated for 1 h at 55°C.

#### Human genomic DNA

MRC-5 cells were pre-treated with 1 mg/ml collagenase prior to trypsinisation. Cell were pelled (MRC-5 after trypsin neutralisation, HeLa S3 – after FACS) and resuspended in 10 mM Tris-HCl pH 8.0, 0.5% w/v SDS, 100 mM EDTA supplemented with 20 μg/ml RNase A and 1 mg/ml proteinase K and incubated at 50°C for at least 1 h.

After the RNase A and proteinase K treatment, DNA samples from yeast and mammalian cells were allowed to cool to room temperature and sample volumes were adjusted to 0.5 mL by addition of TE buffer (10 mM Tris pH 8.0, 1 mM EDTA). The samples were mixed with an equal volume of phenol:chloroform:isoamyl alcohol pH 8.0 (25:24:1; Sigma) followed by single or double extraction using an equal volume of chloroform. DNA was then precipitated using isopropanol in the presence of potassium acetate. DNA concentration was measured using the Qubit™ dsDNA HS assay (ThermoFisher).

### ddPCR

Genomic DNA was cut with *EcoRI, BamHI, Hindlll,* or *Kpnl* (NEB) and diluted to 50-150 pg/μl. Each reaction consisted of 0.5-1.5 ng of yeast genomic DNA or 25-75 ng of human genomic DNA, 1x QX200™ ddPCR™ EvaGreen Supermix (Bio-Rad) and primers (225 nM final each). The samples were processed using the QX200™ Droplet Digital™ PCR system and analysed with the QuantaSoft software (Bio-Rad). Probe primers are listed in Supplemental Tables 2 and 3. A more detailed ddPCR protocol can be found in the Supplemental Materials.

### Sort-seq

Extracted genomic DNA samples were fragmented by sonication so that the majority (~95% or more) of DNA fragments were between 50–500 bp, and the mean length between 200–300 bp as confirmed by TapeStation (Agilent). 500 ng of fragmented genomic DNA was used for library construction for Illumina sequencing. Indexed genomic DNA libraries were prepared using a NEBNext Ultra II library prep kit for Illumina without size selection, and NEBNext Multiplex Oligos for Illumina using four cycles of amplification (NEB), followed by two rounds of clean-up using Agencourt AMPure XP beads (Beckman Coulter). Library samples were quantified by qPCR using a NEBNext Library Quant Kit for Illumina (NEB) and a Rotor-Gene real time PCR cycler (Qiagen). Fragment sizes were confirmed by Tapestation (Agilent). Library samples were mixed and diluted to 2.2 pM for single-end deep sequencing by NextSeq 500 using a NextSeq 500/550 High Output v2 kit (80 cycles) (both Illumina) generating 168,000,000 and 238,000,000 reads for G1 and S samples, respectively.

Sequencing reads aligned (using STAR v2.5.3) to a single genomic location (with up to 2 mismatches) on the hg38 human genome assembly (without chrY) were indexed and summed in 50 kb genomic bins using Samtools (version 1.3.1), Bedtools (version 2.26.0) and custom bash scripts (https://github.com/DNAReplicationLab/batrakou2018.git). In the R environment, the ratio between read numbers from replicating and non-replicating samples was calculated for each genomic bin using the following formula: r = (rep/nonRep) * (nonrepSum/repSum), where r is the calculated ratio, rep is number of reads in a single bin from a replicating sample, nonRep is the number of reads in a single bin from a non-replicating sample, repSum is the total number of reads from a replicating sample and nonRepSum is the total number of reads from a non-replicating sample. The resulting ratio was additionally adjusted using a custom function that minimises the sum of ratio values outside of the one to two range (https://github.com/DNAReplicationLab/batrakou2018.git).

### Data analysis and figure generation

#### Flow cytometry

To estimate bulk genome replication in arrest/release experiments, population means of DNA fluorescence signals were extracted for each sample using FlowJo software, normalised to the arrested sample and the dynamic range was fit between 1 and 2 using a linear contrast stretching algorithm. A Boltzman sigmoid function was fit to the data using the *nls* function of the R base package (33).

#### Comparison to sort-seq

Previously published Illumina sort-seq data (1) was smoothed using the cubic spline function of the R base stats package and values from 1 kb windows overlapping with the location of the corresponding ddPCR probes were extracted. In cases where a ddPCR probe spanned two sort-seq windows, an average of the two windows was used. Linear fit between ddPCR concentration and the smoothed sort-seq relative copy number values was performed using the *lm* function of the R base package.

#### Statistical analysis of the resolution of ddPCR

The experiments in Figure 3 and 5 were performed using three technical replicates and analysed using one-way ANOVA (*aov* function) followed by post hoc Tukey HSD comparison (*TukeyHSD* function) of the R base stats package.

#### Curve fitting

Boltzman sigmoid curves were fit to the replication dynamics data in Figure 4B using the *nls* function of the R base stats package.

#### Figure generation

The figures were produced using ggplot2 R package (34) and Xara Photo & Graphic Designer.

## Results

### ddPCR measurement of relative copy number in non-replicating and replicating cells

The advantages of ddPCR prompted us to explore the possibility of using it as a method to measure relative DNA replication timing. Based on available replication timing data for the S. *cerevisiae* genome (1), we selected several unique regions that replicate either early in S phase (close to early efficient origins), mid/late (a late origin) or late (passively replicated regions of the genome). As well as these single-copy probes, we also designed probes to regions that have either two (*TEF1* and *TEF2* genes) or three (mating loci *HML, HMR* and *MAT)* copies per haploid genome (Supplemental Figure 1A). Each probe consisted of primer pairs that amplified DNA fragments ranging in size (86-197 bp) and GC content (33-49%).

Using these probes, we first analysed DNA from a haploid wild type strain arrested in G1 phase with α-factor. The cells should not have been replicating, therefore unique probes should have a similar concentration, corresponding to a relative copy number of 1. Flow cytometry analysis of the DNA content in the arrested cells showed that 95% were in G1 phase of the cell cycle, with only 3% in S phase (Supplemental Figure 1B). Thus, the replication timing of the amplified regions should not affect the copy number analysis by more than 3%. The top panel of Figure 1A shows that all unique probes had a similar concentration, while the *TEF1/TEF2* and *MAT* probes were double and triple the concentration of the unique probes, respectively.

**Figure 1.**
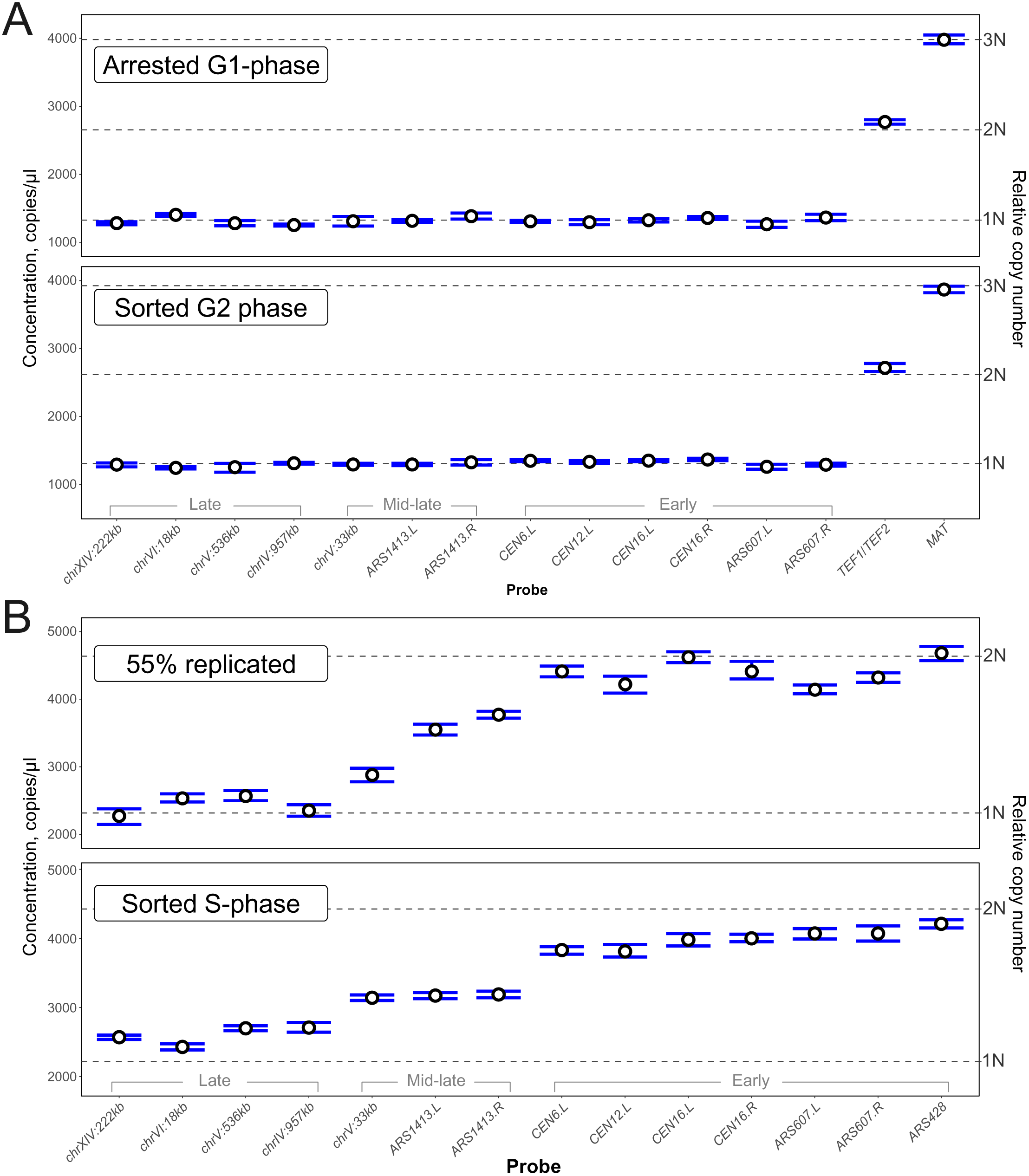
DNA copy number in non-replicating and replicating cells. *A,* DNA copy number in non-replicating yeast cells. *Top,* haploid cells (T7107) were arrested in G1 phase with α-factor and their DNA was analysed by ddPCR. *Bottom,* FACS-enriched G2 phase diploid cells (T9475) were analysed by ddPCR. Indicated probes targeted unique, duplicated or triplicated regions. The left y-axis indicates absolute concentration in the analysed samples. The right y-axis indicates the DNA copy number relative to the mean concentration from all probes (copy-adjusted for non-unique probes). *B,* DNA copy number in replicating cells was analysed by ddPCR. *Top,* DNA from synchronised haploid cells (T7107), in mid-S phase. *Bottom,* DNA from FACS-enriched S phase diploid cells (T9475). ddPCR was performed with probes targeting late, mid-late and early replicating regions. Error bars are 95% CI based on 3 technical replicates.

Fluorescence-activated cell sorting (FACS) is a valuable method for DNA replication studies because it allows the enrichment of cells based on DNA content, and is often used as an alternative to cell cycle synchronisation for species that are difficult to synchronise. As a proof of concept, we sorted G2 phase cells from an asynchronous culture of diploid wild type cells. Flow cytometry analysis of the sorted cells showed that 94% of cells were in G2, with about 3% of cells in S phase (Supplemental Figure 1B), therefore the difference in concentration between an early and late replicating region should not exceed 3%. DNA extracted from these cells was subjected to a similar analysis of integer copy number variation analogous to the α-factor arrested haploid sample. As with the arrested sample, DNA from the G2 sorted sample showed the discrete distribution of unique and non-unique probes that follow a 1:2:3 ratio (Figure 1A, bottom panel).

Having confirmed appropriate probe behaviour using DNA from non-replicating cells, we then analysed DNA from replicating cells. A haploid S. *cerevisiae* culture was arrested with α-factor and released into S phase, with samples taken at intervals for DNA content analysis and DNA extraction. DNA content-based flow cytometry confirmed arrest of cells in G1 phase and synchronous progression through S phase upon release from α-factor (Supplemental Figure 1C). The kinetics of bulk DNA replication in the cell population was determined based on the median DNA content of the cells at each time point (Supplemental Figure 1D). A mid S phase sample (55% genome replication,

35 min after release) was selected for ddPCR using the unique probes (Figure 1B, top panel). As expected, the concentration of probes targeting early-replicating regions was double that of late probes, with the late-firing origin ARS1413-proximal probes having intermediate values. DNA from S phase cells sorted from an asynchronous culture of a wild type diploid strain showed a similar distribution of probe concentrations (Figure 1B, bottom panel) with a correlation coefficient of 0.91 between the synchronised and sorted cells (Supplemental Figure 2). These results suggest that ddPCR can be used as a tool to determine locus-specific DNA copy number as a proxy for relative DNA replication timing.

### Comparison of ddPCR to high-throughput sequencing

High-throughput sequencing has become the *de facto* standard to determine genome replication dynamics in various organisms, including S. *cerevisiae.* It can be used to determine the relative DNA copy number genome-wide as a proxy for DNA replication timing. The temporal and spatial resolution of this method depends on genome coverage – the more reads per bin, the smaller the standard error. At around 1000 reads mapped per 1 kb bin of the S. *cerevisiae* genome, the coefficient of variation (CV) is ~5% (1). Sort-seq and repli-seq analyses of large mammalian genomes typically use larger bins of 10-50 kb in order to increase sequencing depth per bin, thereby retaining temporal resolution at the expense of spatial resolution (8, 14, 16). Similar to the high-throughput sequencing-based sort-seq, ddPCR can determine relative DNA copy number. Errors in ddPCR come from subsampling and partitioning; within a range of 0.11-5.73 copies/droplet (i.e. over a ~50-fold range of concentration) the CV of the absolute DNA concentration calculation is less than 2.5% (31). These errors will limit the temporal resolution, while the spatial resolution is limited by the minimal amplicon length (60 bp) and primer design constraints. Thus, ddPCR is capable of precise relative copy number determination with high spatial resolution that is currently not practical for the high-throughput sequencing-based methods.

Therefore, we decided to compare the ddPCR-determined concentration values to deep sequencing-derived relative copy number values (1). The bulk DNA replication value of the time course sample from Figure 1B closely matched bulk DNA replication from a published deep sequencing time course experiment (55% replicated sample corresponding to 45 min in (1)). Therefore, we extracted HTS-derived normalised relative copy number values from 1 kb windows that overlap with the ddPCR probes and asked how well they correspond to the ddPCR-determined concentrations (Figure 2A). Inherent variations in culture conditions and sampling make it challenging to have highly comparable samples between biological replicates of arrest/release experiments. Nevertheless, a linear fit between the sequencing- and ddPCR-determined relative copy number measurements had an adjusted coefficient of determination (R^2^) of 0.94, demonstrating good correlation between the two methods.

**Figure 2.**
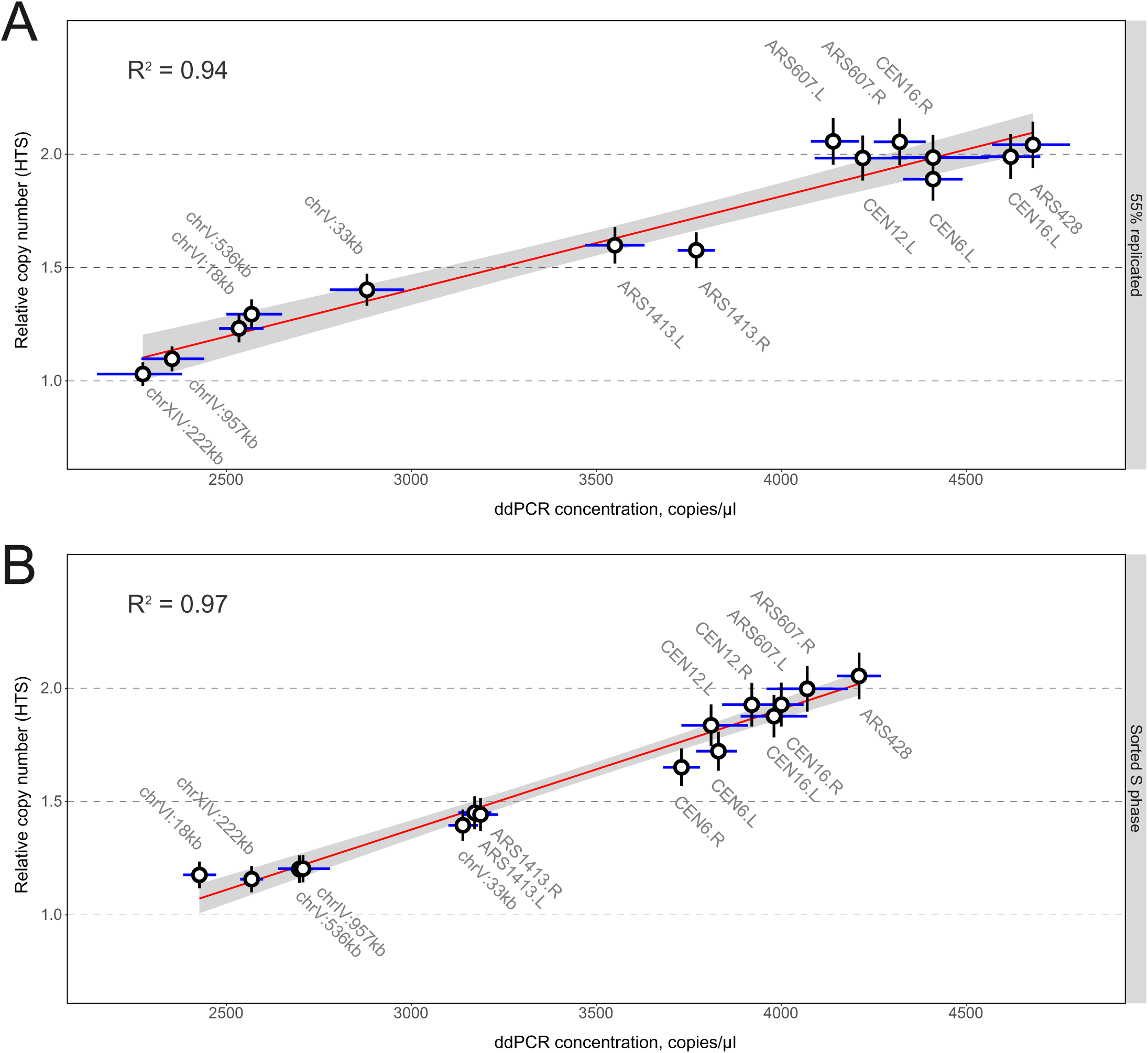
Comparison of ddPCR and HTS-based methods. *A,* linear fit between absolute DNA concentrations measured by ddPCR and HTS (1 from cell-cycle synchronised S phase haploid cells (T7107) with ~55% bulk genome replication. *B,* linear fit between absolute DNA concentration measured by ddPCR and HTS (1 from FACS-enriched S phase diploid cells (T9475). Error bars, HTS: estimated 5% coefficient of variation; ddPCR: 95% CI based on 3 technical replicates.

Consistent gating in FACS can offer more reproducibility and, therefore, more comparable S phase samples. We therefore performed a linear fit between ddPCR- (Figure 1B, bottom panel) and HTS-derived (1) values from FACS-enriched samples (Figure 2B). The linear fit resulted in a R^2^ of 0.97, indicating good agreement between these two methods of assessing relative DNA copy number.

### High resolution measurement of replication time in yeast

To accurately determine replication timing in S phase cells, the resolution of detecting copy number variation should be sufficiently high to be able to distinguish between multiple values on the continuous scale between one and two. The resolution of a similar method, qPCR, depends on the number of technical replicates. Four qPCR replicates allow discrimination between a copy number of 1 and 2 at 95% confidence, with the least stringent Power test, while 17-40 qPCR replicates are required to distinguish between 4 and 5 copies, corresponding to the difference between 1 and 1.2 on the relative copy number scale (26). We tested the resolution power of ddPCR by designing adjacent probes every 2 kb across an approximately 45 kb region of yeast chromosome 4 that previously has been shown to span relative copy numbers between two (near *ARS428*) and one (proximal late replicating region) (1).

The 55%-replicated sample (from Figure 1B) was analysed by ddPCR using a panel of probes spanning the *chrIV* 913-958 kb region. Figure 3 shows the ddPCR-determined relative DNA concentration and equivalent high-throughput sequencing-derived relative copy number values (R^2^=0.91). To analyse the significance of the difference between adjacent ddPCR probes, one-way ANOVA followed by post hoc Tukey HSD was performed on the data from three technical replicates. Eight statistically different states were detected from the chosen set of primers (p<0.05), with 0.07 being the smallest significant detected difference in relative copy number. Therefore, on a scale from one to two, up to 14 ((2-1)/0.07) significantly different states can theoretically be detected by ddPCR. As a result, the resolution of ddPCR is over an order of magnitude higher than that of real time PCR when using three technical replicates.

**Figure 3.**
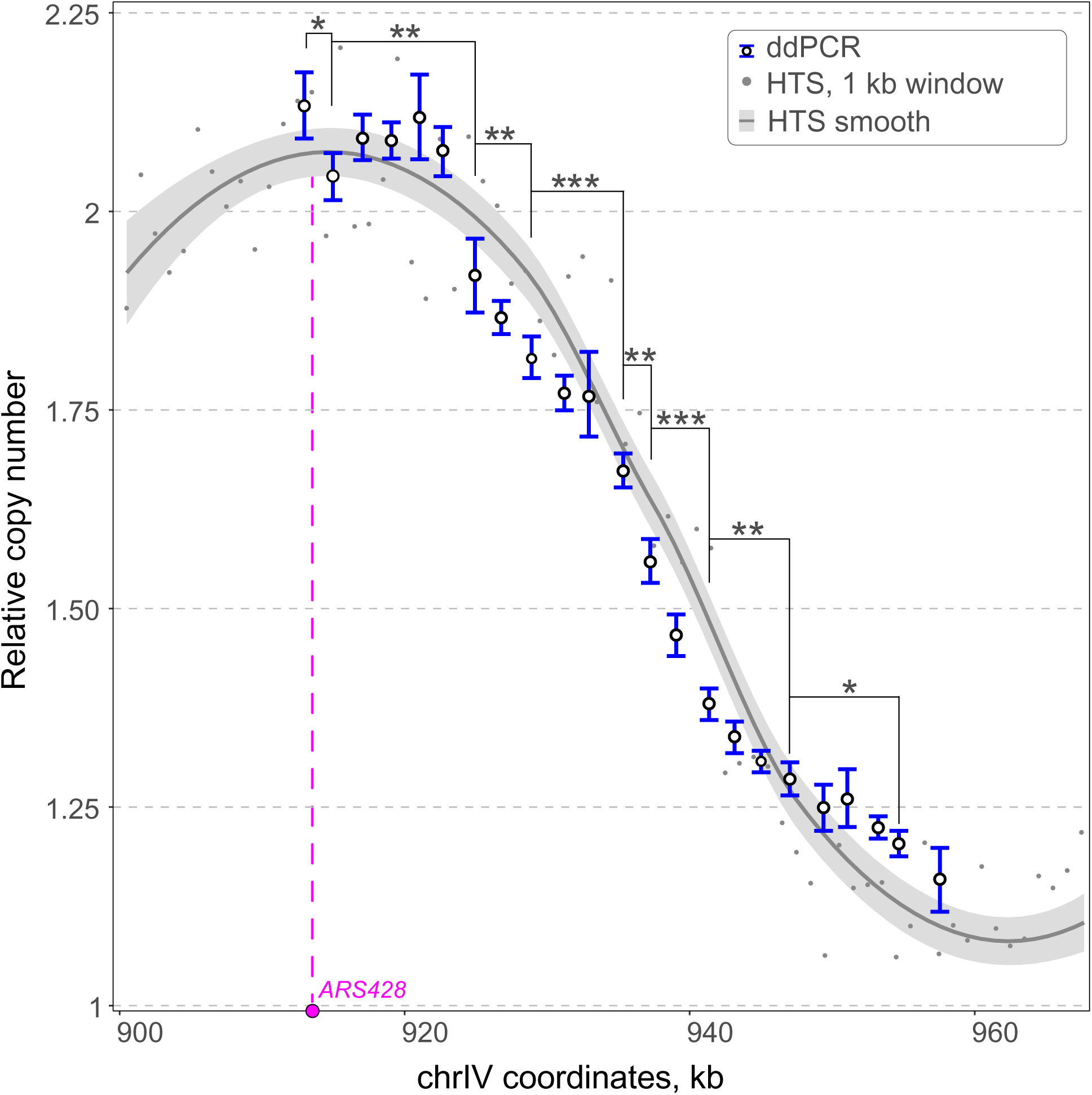
Resolution of ddPCR for measurement of DNA replication timing. Comparison of relative DNA concentrations measured by ddPCR and HTS (1 for the genomic region including *ARS428,* as indicated on the x-axis. Both datasets are from cell-cycle synchronised S phase haploid cells (T7107) with ~55% bulk genome replication. The HTS data is shown with and without smoothing. The ddPCR data error bars are 95% CI based on technical replicates. HTS data is smoothed using LOESS local regression, shown with 0.95 confidence interval. Asterisks denote significance in ANOVA followed by Tukey HSD test (* p-value ≤ 0.05, ** p-value ≤ 0.01, *** p-value ≤ 0.001).

### Inter-sample comparison of replication timing using ddPCR

Often it is necessary to compare the replication timing of a DNA region between two or more independent samples and that brings the challenge of inter-sample normalisation. Ideally, the samples to be compared need to have exactly the same amount of DNA (in the case of equal S phase populations, i.e. DNA from the similarly sorted cells) or adjusted according to percentage of the bulk genome replication. Traditionally-used methods of DNA quantification are not sufficiently quantitative for the task. Absorbance-based methods require very pure samples for quantification of DNA to be accurate, while the SYBR Green-based approach (as used in the Qubit™ system) can still lead to an error of 20% in the estimation of the DNA concentration, and it is less accurate if samples contain single-stranded DNA.

To address this issue, we used the commonly applied approach of control probes. For sample normalisation, a control probe must remain at a constant concentration between the different samples. This is straightforward for pooled S phase samples, for example as obtained by cell sorting. However, it is more challenging for DNA replication samples from a synchronised arrest-release experiment, since every allele replicates (increases in concentration) at some point during the time course. Therefore, we tested whether time points in early S phase could be normalised to a late replicated region that has not started to replicate (constant concentration). Later time points could then be normalised to an early-replicating region that has completed DNA replication (constant concentration). In order to assess all time points, the early region must be completely replicated before the onset of the replication of the late region. To this end, we chose a probe targeting a late-replicating region on chromosome 4 (*chrIV:966kb*) and a probe proximal to an early-firing origin *(ARS607)* (1). We performed an arrest-release experiment (Supplemental Figure 3) and analysed the concentration of these probes by ddPCR. Figure 4A shows the concentration ratio of these control probes as a function of time after release from the G1 arrest. As expected, in the arrested G1 cells the ratio is close to 1 indicating that neither of the loci tested have been replicated. In the first half of S phase, the ratio progressively approaches 2, as the early locus is being replicated, while the late locus is not. At ~50% bulk replication (44 min after release), we observe a maximum ratio which is close to 2. The observed ratio of ~1.8 is, most likely, lower than two due to imperfect synchronisation and in a biological replicate it approached 2 (Supplemental Figure 4). Therefore, the early locus is replicated in almost all the cells prior to the late locus starting to replicate. In the second half of S phase, the ratio decreases as the late locus is replicated in a progressively larger proportion of cells, and in G2 cells both loci are fully replicated, thus giving a ratio of 1. This confirms that it is possible to use these probes for dual normalisation in a synchronous S phase experiment, with the late probe used until 44 min, at which point the early probe can be used (adjusted for ploidy).

**Figure 4.**
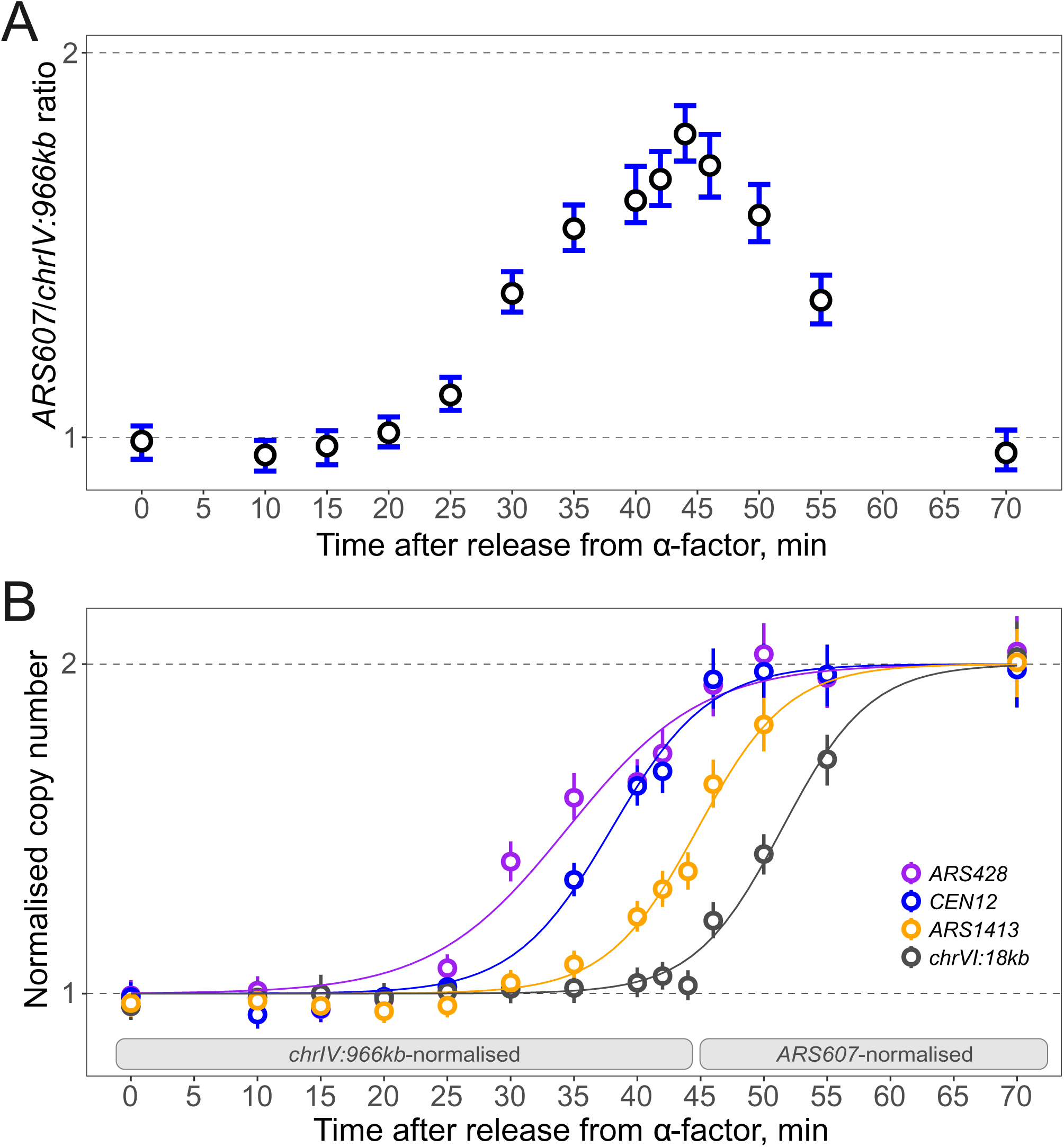
DNA replication dynamics during a synchronised cell cycle time course. *A,* DNA concentration, determined by ddPCR, of an early *(ARS607.L2)* relative to a late replicating probe *(chrIV:966kb)* through a synchronised cell cycle. Haploid cells (T7107) were arrested in G1 phase and synchronously released into S phase and samples taken at the times indicated on the x-axis. *B,* Normalised copy number of early *(ARS428),* mid-early *(CEN12.L),* mid-late *(ARS1413.L)* and late *(chrVI:18kb)* replicating probes. Samples from early S phase (up to and including the 44 min sample) were normalised to the late *chrIV:966kb* probe; samples from late S phase (46 min onwards) were normalised to the early replicating *ARS607.L2* probe, adjusted for copy number. Error bars are 95% CI based on 2 technical replicates. Curve fits were generated by fitting to a sigmoidal Boltzmann function.

As a proof of principle, we determined replication kinetics for early *(ARS428),* mid/early *(CEN12.L),* mid/late *(ARS1413.L)* and late *(chrVI:18kb)* probes, applying the dual normalisation (Figure 4B). The loci replicated in the order *ARS428, CEN12, ARS1413* and *chrVI:18kb* as anticipated from previous studies and with similar relative kinetics (1, 18, 35, 36). Therefore, ddPCR can be used to determine locus replication timing across a time course experiment.

### Allele-specific replication timing

Small changes in DNA sequence can lead to large changes in replication timing. For example, replacing just a few nucleotides within the ORC-binding site can lead to complete origin inactivation in S. *cerevisiae* (37). Diploid heterozygotes, where one allele is mutated while the other one is wild type, present a rare example of an internally-controlled system. Here, the defined change is contained within the same cell as the control, and both are subjected to the same conditions. PCR-based methods can be used to distinguish even a single nucleotide polymorphism (38). As a proof of concept, we used a diploid heterozygous strain that contains three inactivated origins of replication *ARS606, ARS731.5* (also known as *ARS737)* and *ARS1021* (previously *ARS121),* as previously described (37). The origins are inactivated by point mutations in their respective ORC-binding sites, changing the critical AT-rich region to the *XhoI* restriction enzyme recognition site (CTCGAG), thus the alleles are only different by up to 6 nucleotides. We designed and confirmed primer pairs with specificity to either the wild type allele or the mutant allele. S phase cells were sorted from an asynchronous culture of the heterozygous diploid and the extracted DNA was analysed by ddPCR using the allele-specific probes, as well as pan-allelic controls (early *ARS607*-proximal and late *chrXIV:222kb* probes). As expected, the DNA concentration from the allele-specific probes was lower than the probes that amplify from both alleles (Figure 5). Furthermore, there was a clear difference in concentration between wild type origins and inactivated origins, with *ARS737* and *ARS1021* showing the strongest effect. The smaller difference at *ARS606* may be explained by a proximal early origin *(ARS607)* that passively replicates the *ars606* locus. Therefore, ddPCR has the power to distinguish alleles with minimal differences in sequence, consistent with a previous report using asynchronously growing cells (8).

**Figure 5.**
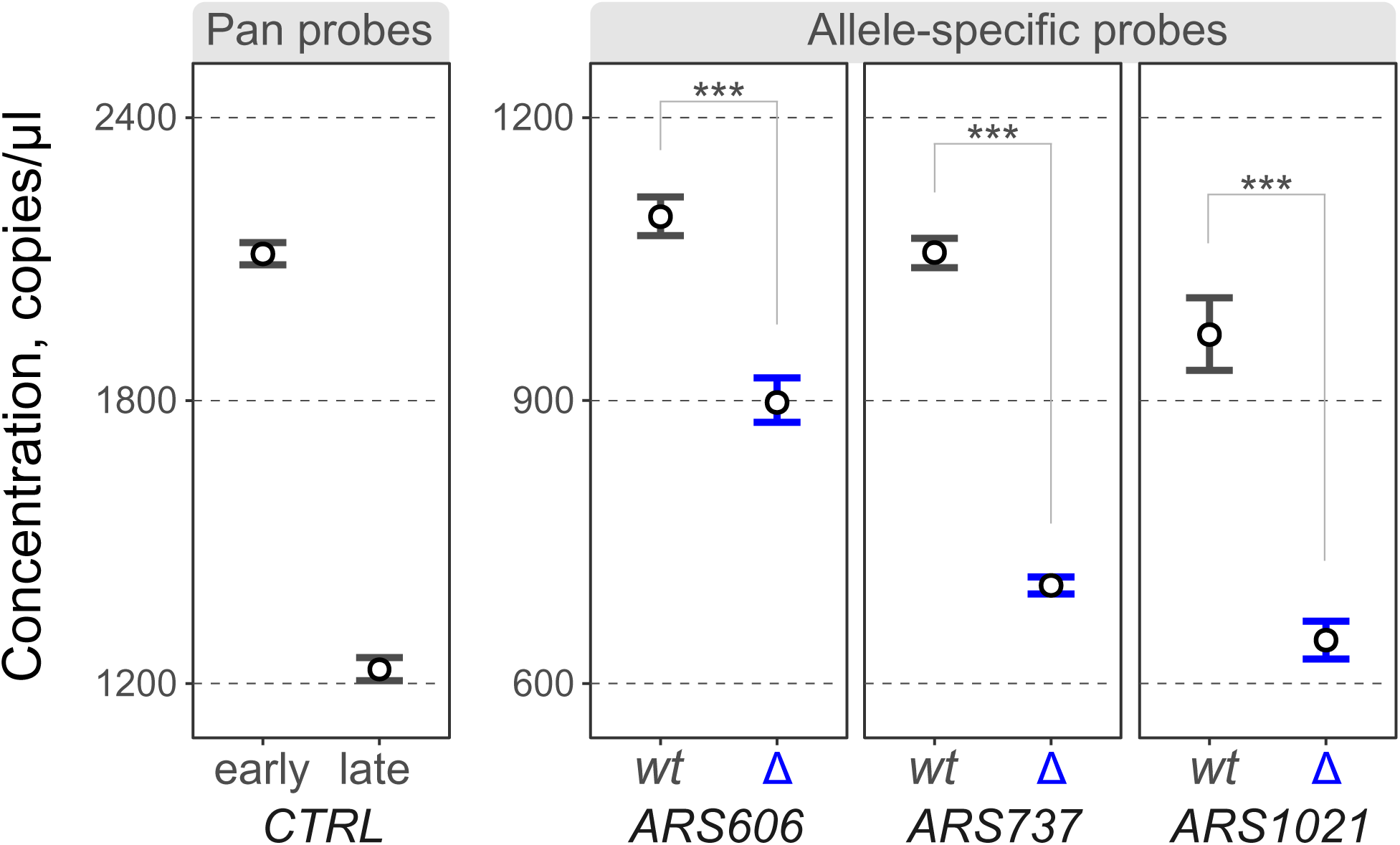
Allele-specific DNA replication timing. S phase cells were FACS-enriched from an asynchronous population of diploid cells (MHY230). The cells are heterozygous for three origins *(ARS606, ARS737* and *ARS1021)* due to one allele of each origin containing an origin-inactivating mutation. Extracted DNA was analysed using allele-specific or pan-allelic probes. Early control, *ARS607.L2* probe; late control, *chrXIV:222kb* probe. Each mutated allele (labelled ∆) shows lower copy number than its wild type (labelled wt) counterpart corresponding to the delay in replication time. Note that the allele-specific probes (right panel) are on a different y scale than the pan-allelic probes (left panel). The error bars are 95% CI based on 3 technical replicates. Asterisks denote P-value ≤ 0.001.

### Detection of replication timing transition zones in human cells

Lack of suitable locus-specific methods to determine replication timing is especially detrimental to studies in organisms with large genomes, such as mammalian cells. Available locus-specific methods are elaborate and of low resolution (39). Alternative genome-wide replication timing profiling, aided by HTS, either requires a large read number in case of sort-seq, or elaborate sample processing during the repli-seq procedure. Therefore, we tested whether ddPCR could be used as a rapid and cost-effective way to analyse the relative replication timing in locus-specific mammalian studies. To this end, we first designed sex chromosome-specific probes, as well as probes targeting autosomal and duplicated autosomal regions of the human genome (40). We tested these probes on DNA from the male MRC-5 cell line arrested in G1 phase (Figure 6A, Supplemental Figure 5). As expected, autosomal probes *(TOP1, MYC1, chr18:4.6Mb)* and a probe targeting both sex chromosomes *(PLCXD1)* were twice the concentration of unique sex chromosome probes *(KAL1, XIST, SRY,*

**Figure 6.**
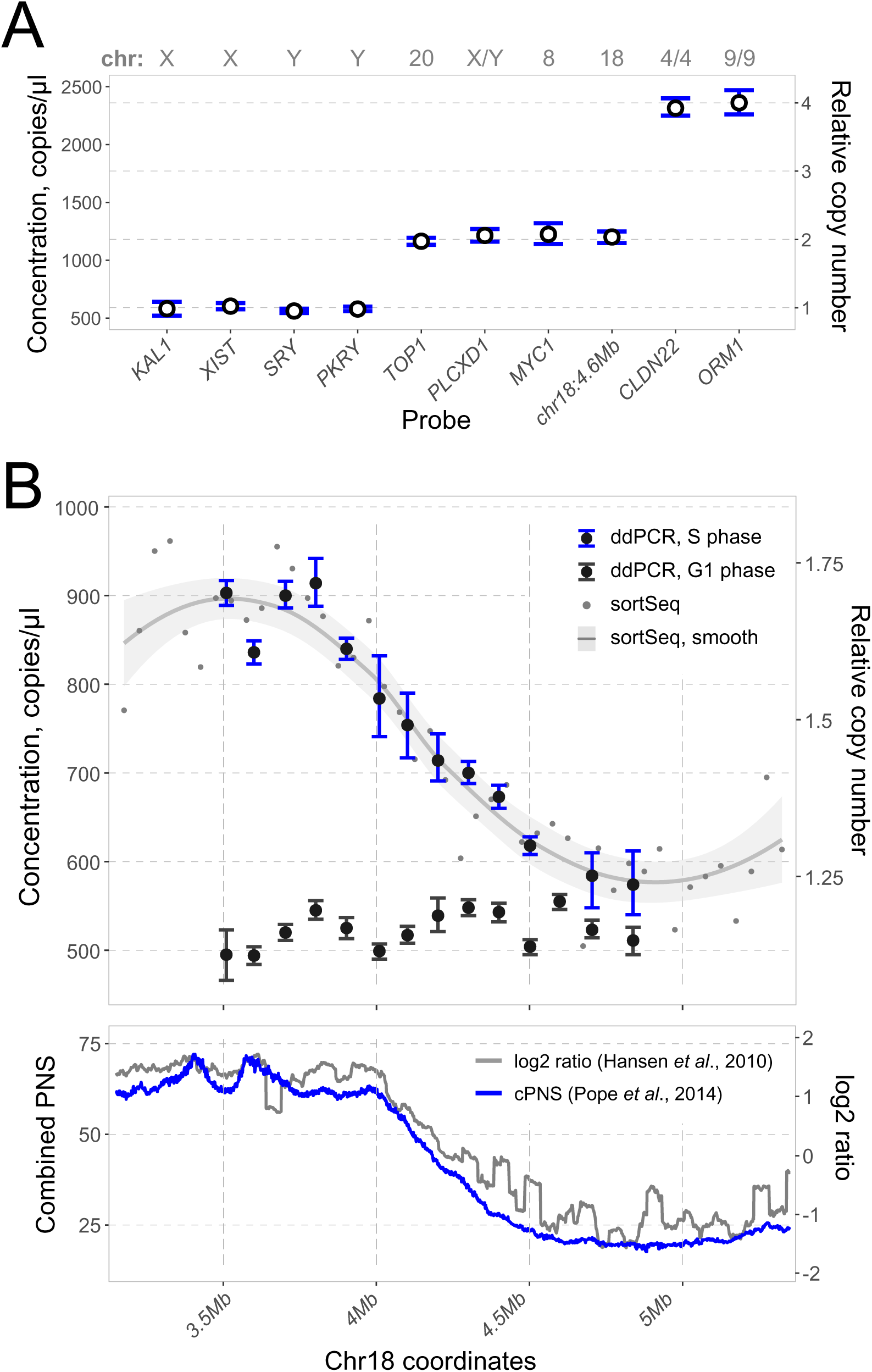
Application of ddPCR to determine relative DNA replication time in human cells. *A,* DNA copy number analysis in non-replicating human cells. DNA from male MRC-5 cells arrested in G1 phase of the cell cycle was analysed by ddPCR with probes targeting sex chromosomes, autosomal chromosomes and duplicated genes. Relative copy number was calculated as a mean concentration of all probes adjusted by copy number. *B*, DNA concentration analysed by ddPCR across a known replication timing transition zone on chromosome 18. *Top:* non-replicating control from FACS-enriched G1 HeLa cells, as well as FACS-enriched S phase HeLa cells analysed by ddPCR and sortSeq. Note that G1 and S phase samples, while coming from the same initial population of cells, are independent DNA samples and their absolute concentration cannot be compared. *Bottom:* repli-seq data of the corresponding genomic region from (14, 41). cPNS - combined percentage-normalised signal.

*PKRY).* In addition, the concentration of duplicated autosomal probes *(CLDN22* and *ORM1)* was four times the concentration of unique sex chromosome probes.

To test the dynamic range of ddPCR in sorted mammalian cells, we analysed available HeLa cells repli-seq data (41) and designed probes that span a ~1.5 Mb region on chromosome 18 that contains a sharp change in replication timing. Using only DNA content, we sorted cells from S and G1 phases, extracted the DNA and analysed DNA copy number by ddPCR and sort-seq. ddPCR analysis of the G1 sample showed the probed loci to be at similar concentrations (Figure 6B and Supplemental Figure 6, top panels), while the concentrations of the loci in the S phase sample closely matched relative copy number determined by sort-seq (R^2^ of the linear fit is 0.94). The data from both relative copy number methods resembled the data from the nascent-strand repli-seq approach (Figure 6B, bottom panel) (14, 41). We also tested a region in the 11q chromosome locus in Jurkat cells, which has been shown to have a sharp transition from early to late replication by a locus-specific nascent DNA-based method in a closely related THP-1 cell line (39). Similar to the HeLa cell transition zone, probes amplified from the Jurkat S phase sample had concentrations that corresponded to the expected transition between early and late replication, while their concentration was uniform in the G1 control sample (Supplemental Figure 7). Therefore, ddPCR is able to detect replication timing transition zones in cultured mammalian cells and offers a relative copy number measurement that may be used as an alternative to HTS-based methods to determine replication timing of defined loci in organisms independent of genome size.

### Detection of centromere DNA replication differences by ddPCR

We next aimed to compare replication timing in multiple biological samples by ddPCR. We focused on replication of centromeric DNA in S. *cerevisiae,* since we have previously shown that Dbf4 enrichment at kinetochores leads to early activation of centromere-proximal origins (42). C-terminally tagged Dbf4 is no longer associated with the kinetochore and replication of the centromeric DNA is delayed in this mutant. We tested the replication timing of several centromeres (each with two proximal probes less than 10 kb apart), both in wild type and Dbf4-9myc tagged cells. The tested centromeres included ones previously shown to be strongly affected by Dbf4 tagging *(CEN9, CEN12, CEN16)* and unaffected by tagging *(CEN2, CEN4, CEN6)* (42). S phase cells from asynchronous cultures were sorted and DNA was extracted for analysis using ddPCR. The two samples had different dynamic ranges, most likely due to variation in the sorted fractions (Supplemental Figure 8A). To account for this, we used replication index as a way to compare the replication timing of the probes between the samples. Replication index has been extensively used to analyse density transfer experiments (43). It compares median replication time (T^rep^) of probes on a scale from 0 to 1, with 0 being the T^rep^ of a control early probe and 1 the T^rep^ of a control late probe. In an analogous manner, we used the concentration of ARS607-proximal and *chrXIV:222kb* probes to normalise other probes. Figure 7A shows the replication indices for the tested centromeres, as well as independent early *(ARS428-* proximal) and late-replicating *(chrIV:966kb)* controls. Similar to the HTS-derived data, centromeres 9, 12 and 16 had their replication time delayed in the Dbf4-9myc mutant, while centromeres 2, 4 and 6 were unaffected. Thus, ddPCR was able to confirm the effect of Dbf4 C-terminal tagging on centromere replication timing.

**Figure 7.**
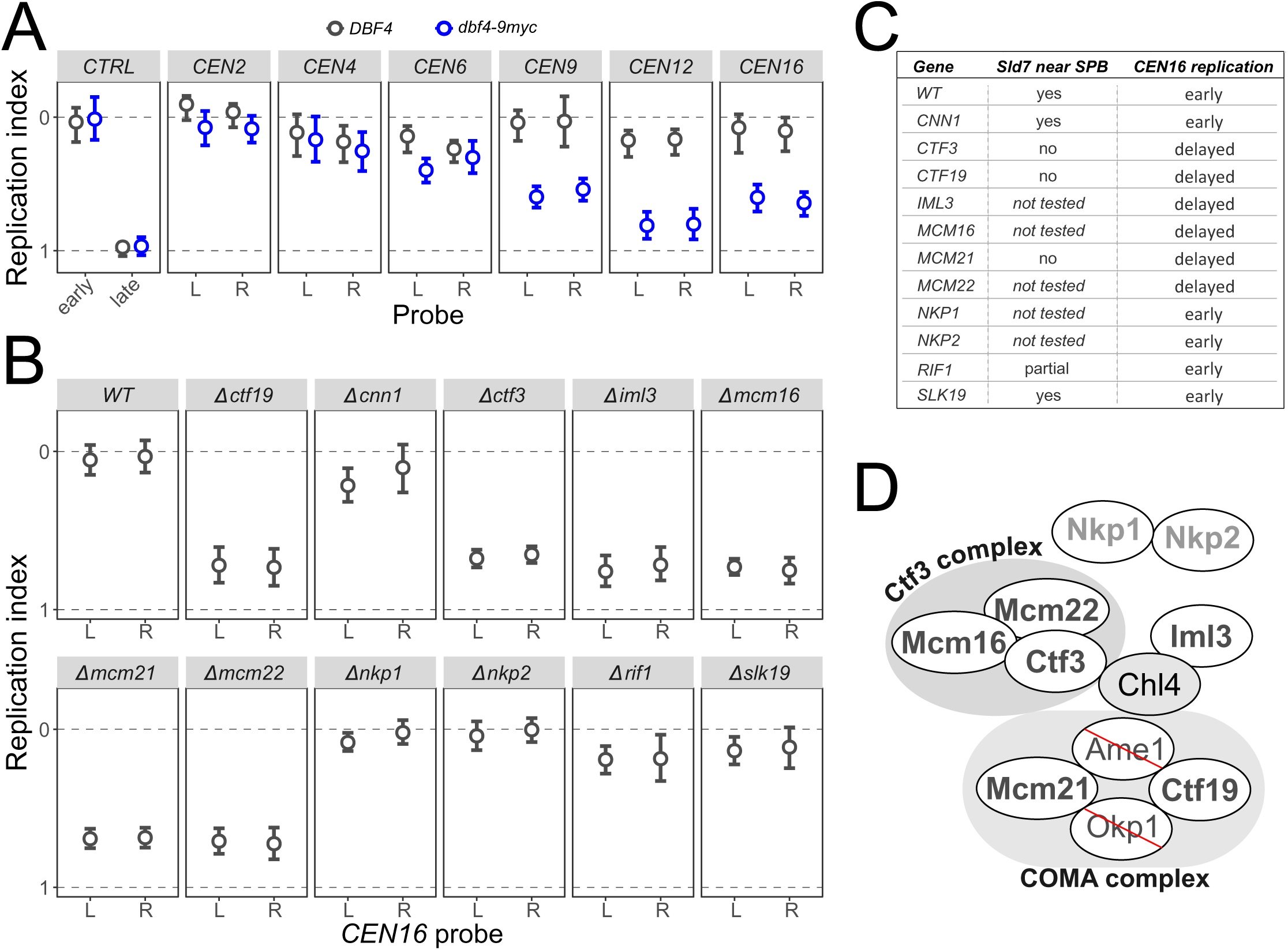
Identification of *trans*-acting factors regulating centromere replication timing. *A,* Replication of DNA at centromeres is delayed in a *dbf4-9myc* mutant, as previously reported (42). Replication timing of 6 centromeres was analysed by calculating replication indices (where zero and one are defined as the S phase DNA copy number of *ARS607.L2* and *chrXIV:222kb,* respectively). Additionally, control early *(ARS428)* and late *(chrIV:966kb)* probes are shown. *B,* Application of ddPCR to rapidly screen for *trans*-acting factors that affect DNA replication timing. Replication timing of two *CEN16* proximal probes (left and right) was analysed in wild type cells or cells with the indicated gene deletions. *C*, Table summarising results in B and comparing them to a previously reported screen (42 for Sld7 co-localisation with the spindle pole body (SPB). *D,* Schematic representation of the Ctf19 kinetochore sub-complex (modified from (51) indicating components that affect centromere replication timing. In bold – tested components; in dark grey – components whose deletion affected replication timing, in light grey – components that did not affect replication timing; crossed – essential components that were not tested.

In addition to the Dbf4 mutant, it has been shown that a *CTF19* deletion mutant also has a similar effect on centromere replication timing (42). Previously, Sld7 co-localisation near the spindle pole body was used as a read-out to screen for the components of the kinetochore required to localise replication factors to centromeres (42). Here, we used ddPCR to screen for delay to *CEN16* replication timing in deletion mutants of components of the Ctf19 complex. In addition, we screened deletion mutants of *CNN1, SLK19* (non-Ctf19 kinetochore components) and *RIF1,* a negative regulator of DNA replication (44). As above, results are expressed as replication index (Figure 7B, Supplemental Figure 8B), and are summarised in Figure 7C & D. We find that all mutants that lost Sld7 localisation also showed a delay in *CEN16* replication timing. Additionally, deletion of several genes that were not tested in the Sld7 localisation screen also showed delayed *CEN16* replication timing. These included *MCM16* and *MCM22* (encoding components of Ctf3 subcomplex), as well as *IML3* (Chl4 subcomplex). In contrast, *NKP1* and *NKP2* (NKP subcomplex) deletions did not affect replication timing. This targeted screen demonstrates the throughput of the ddPCR methodology and its ability to rapidly identify *trans*-acting factors affecting DNA replication timing of a specific locus.

## DISCUSSION

Measurement of DNA replication timing during S phase can be achieved via measurement of relative DNA copy number. This approach can be used with DNA extracted from either an asynchronous cell population (for example, marker frequency analysis in bacteria) or from S phase cells enriched for by synchronisation or using FACS. This has recently been exploited in HTS-based methods to generate genome-wide replication profiles in various species (6, 8, 12, 45–47). However, current locus-specific approaches are technically challenging, time consuming, expensive and/or of lower resolution. These restrictions have limited the ability to screen for *trans-* or cis-acting factors involved in locus-specific regulation of DNA replication timing. Here, we demonstrate that the high-resolution of ddPCR, as a measure of DNA copy number, allows accurate determination of relative DNA replication timing. While ddPCR gives data comparable to those from HTS, it offers complementary applications particularly for locus-or allele-specific analyses, for screening panels of mutants, and for applications in organisms with large genomes.

Recently ddPCR has been used to quantify the number of rDNA repeats in S. *cerevisiae* within a range of 20–1000 copies per genome with 5-10% technical error (48). Our results suggest similar levels of technical error in measurements within the range of one to two during DNA replication (Figure 3). The technical errors in ddPCR come mainly from sub-sampling and droplet partitioning, where the former prevails in low abundance samples while the latter in high copy number samples. Therefore, ddPCR has the greatest certainty at intermediate concentrations of target DNA. Concentrations within the range of 110 – 5730 copies/μl have CVs less than 2.5%; a >50-fold dynamic range. The naturally limited two-fold dynamic range of DNA replication allows the use of the same dilution for both early and late replicating probes within the range of minimal technical error, corresponding to concentrations between 1000 and 2000 copies/μl and a CV of ~1%. This small measurement error allows precise comparison of probe concentrations within a single biological sample and, when comparing replication timing between samples with large biological variation, it contributes very little to the total uncertainty. While we have not tested other digital PCR platforms, it is likely that our findings will be applicable to other digital PCR techniques.

The analysed cell population will have an impact on the dynamic range of the relative copy number with variation ranging from 1 to 1 + [proportion of cells in S phase]. Thus, if an asynchronous population of cells is analysed, of which 20% of cells are in S phase, the dynamic range would be 1 to 1.2. By comparison, an S phase cell population from the most efficient cell cycle synchronisation will have a dynamic range approaching 2 where there are loci that are fully replicated before other loci commence replication (Figure 4A). Similarly, FACS aims to isolate S phase cells from asynchronous population. However, Gaussian dispersion of DNA dyes used during FACS causes overlapping signal at the phase boundaries (G1/S and S/G2) leading to underrepresentation of cells from the earliest and latest stages of S phase and contaminating the sample with non-replicating G1 and G2 cells. This contributes to the observed reduction in the dynamic range of sorted S phase samples when compared to a mid S phase sample from a well-synchronised yeast cell population (Figures 1 and 2, Supplemental Figure 2).

The replication time of a DNA locus can be influenced by its sequence. Thus, heterozygous alleles in a diploid organism may have different replication times (8). Approaches that are able to distinguish replication timing between the two alleles have two main benefits. First, the two alleles are in the same cell and this provides an identical environment for both the mutated (experimental) and the control wild type alleles. Second, there is no need for inter-sample normalisation using extra probes, which would lead to amplification of experimental error. The effect of a mutation on replication timing can simply be expressed as a ratio to the control wild type allele. In PCR, careful primer design enables highly specific amplification which can distinguish DNA molecules that differ by only one nucleotide. The positional requirement of the distinguishing nucleotide greatly limits choice in primer design. For accurate quantification, real-time quantitative PCR requires primer efficiency to be close to two (1.8-2.1) and thus, occasionally, may be incompatible with assays where primer position is restricted. In comparison, ddPCR uses end point amplification allowing low primer efficiency to be counteracted by an increased number of amplification cycles. As a result, ddPCR has the power to differentiate between the replication timing of nearly identical alleles in a diploid/polyploid organism (Figure 5).

In comparison to HTS-based approaches, ddPCR has a much lower input DNA requirement per sample. In the case of S. *cerevisiae,* to be within the smallest error range, we have used ~0.5-1 ng of DNA per reaction. Typically, it took ~10 minutes to sort sufficient numbers of haploid cells to give enough DNA for 20-60 reactions. This is dramatically less than the ~2 hours of FACS required per sample when subsequent measurements are via HTS. This advantage, coupled with a comparatively low cost per ddPCR reaction and fast turnaround time, allowed us to use ddPCR to screen candidate mutations for an effect on centromere replication timing. As a proof of principle, we were able to screen for changes in DNA replication timing at centromere 16 in twelve deletion strains in under 2 weeks, including time for strain recovery and culture growth. We found that centromere 16 replication timing is delayed in strains lacking *CTF19, MCM21, CTF3, MCM16, MCM22* and *IML3.* A recent report demonstrated direct binding of Dbf4/Cdc7 to Ctf19/Mcm21 dimers and Ctf3/Mcm16/Mcm22 trimers (49), giving weight to the results presented here. However, the same study showed that Chl4/Iml3 dimers did not interact *in vitro* with Dbf4/Cdc7, while we have demonstrated that *IML3* deletion delays *CEN16* replication timing. This apparent discrepancy could be explained in at least three ways. First, deletion of *IML3* may affect overall kinetochore structure and thus reduce Dbf4/Cdc7 binding indirectly. Given that deletion of a single component of either Ctf19 or Ctf3 subcomplexes is sufficient to delay *CEN16* replication timing, it is likely that Dbf4/Cdc7 binding requires an intact kinetochore. Second, it is possible that the *in vitro* assay does not fully recapitulate events *in vivo,* for example, due to a lack of post-translational modifications. Finally, the Chl4/Iml3 subcomplex may be required to advance centromeric DNA replication timing in a second step downstream of Dbf4/Cdc7 recruitment to the kinetochore.

The locus-specificity of ddPCR also allows it to be practical even in organisms with large genome sizes. With HTS-based methods of relative DNA copy number determination there is a trade-off between the cost of sequencing in order to achieve high genome coverage to maintain the same error rate and lower spatial resolution. For example, a sort-seq replication profile of the human genome at 1 kb resolution and 5% CV would require 3.5 billion reads per sample, which is impractical even with recent advances in HTS technologies. In contrast, the technical error of ddPCR depends mostly on the concentration of target DNA in the sample, which can be approximated using various DNA quantification methods. As a proof of principle, we have shown that ddPCR is able to quantify relative DNA copy number across two replication timing transition zones in human cell lines. We note that neither of the tested timing transition zones had the dynamic range of the sorted yeast samples. This is, most likely, due to the fact that transformed mammalian cell lines are heterogenous populations with cells containing variable number of chromosomes. This interferes with the DNA content-based cell cycle stage enrichment during FACS. For example, a proportion of cells that appear to be in G1 phase according to the DNA content fluorescence can, indeed, incorporate BrdU – a characteristic property of S phase cells (50).

As part of this study, we have performed sort-seq on sorted HeLa cells. This allowed us to compare the two HTS methods – relative copy number-based sort-seq and nascent strand-based repli-seq, for which we used published data (14). Comparison of these methods, using 50 kb bins, produced a correlation coefficient of 0.6 (Supplemental Figure 9A), while cubic spline smoothed signal comparison had a correlation coefficient of 0.71 (Supplemental Figure 9B). This correlation is likely an underestimate of the comparability of the two methods due to potential heterogeneity in HeLa cell karyotype between different laboratories (bioRxiv: http://dx.doi.org/10.1101/307421). Additionally, both methods have similar dynamic range. We note that the region on chr18, analysed by ddPCR (Figure 6B) represents almost the full dynamic range observed across the whole genome (Supplemental Figure 10). Therefore, it is possible to use the probes from Figure 6B for inter-sample normalisation in human cells, similar to the strategy presented in this paper using yeast (Figure 4).

In conclusion, ddPCR offers a high spatial and temporal resolution approach to rapidly determine locus-specific replication timing in both sorted cells and synchronised cell populations, including in organisms with large genome sizes. The rapid sample turnaround times make this ddPCR approach a valuable addition to current tools for the study of replication dynamics, particularly for screening candidates, for validating samples prior to HTS, and for allele-specific analyses. Coupled with advances in CRISPR/Cas9 genome editing, it will allow rapid screening of *cis-* and *trans*-acting factors that affect DNA replication timing in yeasts, cultured mammalian cells (bioRxiv: https://doi.org/10.1101/285650) and other model systems.

## DATA AVAILABILITY

Sequenced HeLa raw fastq files and processed bed files giving the final calculated relative copy number values are available from the NCBI GEO database (accession number GSE114480). Genomic data for sort-seq and repli-seq (14) described in this study can be visualised via a UCSC genome browser hub (http://genome.ucsc.edu/cgi-bin/hgTracks?hgS_doOtherUser=submit&hgS_otherUserName=Can1002&hgS_otherUserSessionName=hg38).

## ACKNOWLEDGEMENTS

We are grateful to Tomoyuki Tanaka for kindly sharing yeast strains; to Sally Cowley for MRC-5 cells; to Ulrike Gruneberg for HeLa cells; and to Yi-Chun Yeh for Jurkat cells. We thank Nieduszynski group members for helpful discussion and advice; Michal Maj and Carolin Müller for cell sorting; William James for access to the QX200™ Droplet Digital™ PCR system; Rosemary Wilson, Carolin Müller, James Carrington, Fumiko Esashi, Anne Donaldson and Shin-ichiro Hiraga for critical reading of the manuscript.

As part of this study, the HeLa genome was sequenced using short read Illumina sequencing. The reads were mapped to an ensemble reference hg38 genome and no attempt was made to reconstruct the specific genome sequence. Henrietta Lacks, and the HeLa cell line that was established from her tumor cells without her knowledge or consent in 1951, have made significant contributions to scientific progress and advances in human health. We are grateful to Henrietta Lacks, now deceased, and to her surviving family members for their contributions to biomedical research.

